## supplementary file for "Cassini: Streamlined and Scalable Method for in situ profiling of RNA and Protein"

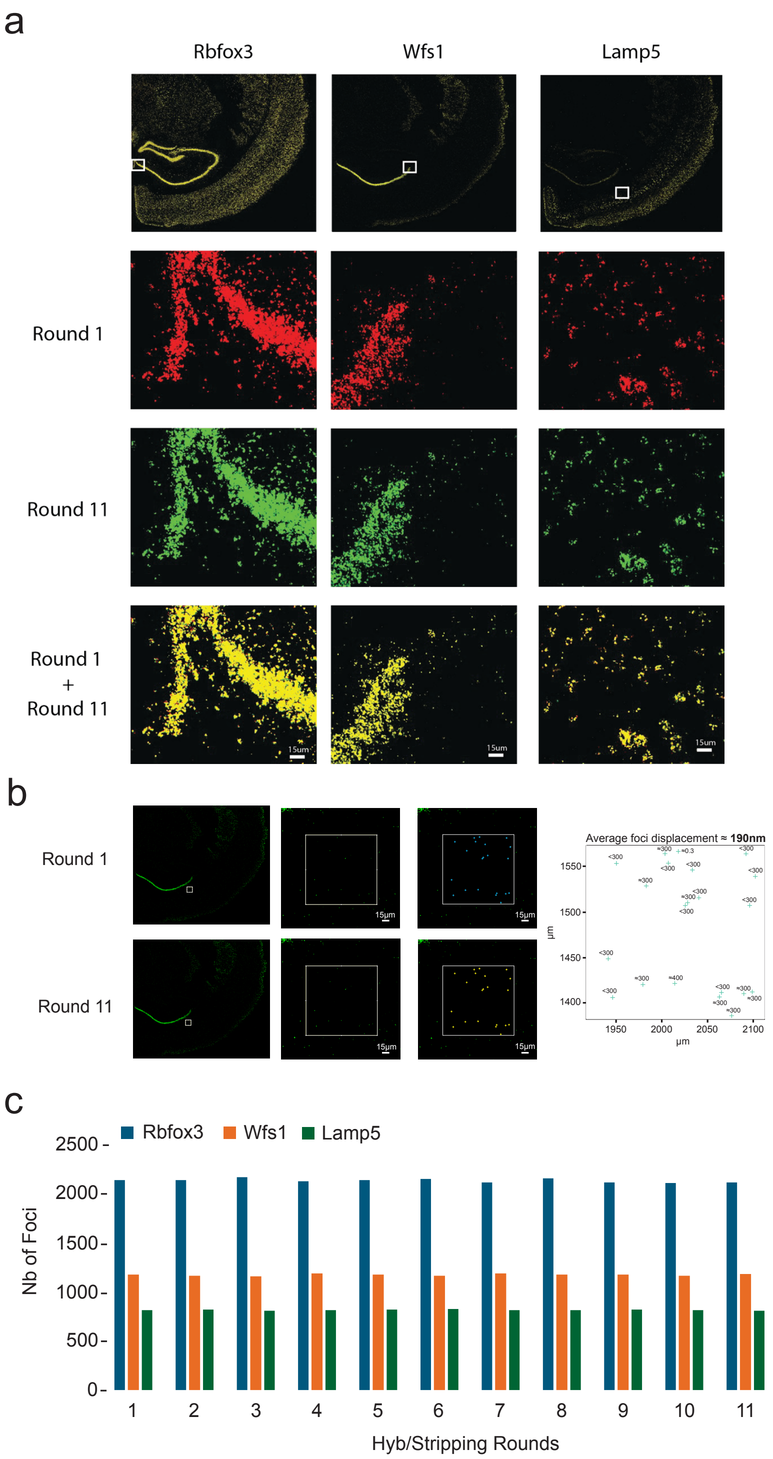


**Supplementary Fig. 1. Foci displacement and sensitivity over Hyb/stripping rounds.** (a-b) Evaluation of displacement of rolling circle amplicons from mRNA targeting padlock probes under multiple rounds of stripping and hybridization. (a) Top row: Full hemisphere showing the merged signals from rounds 1 and 11, with the white square indicating the zoomed areas displayed in the images below. Bottom row: Zoomed-in images of foci from round 1 (red) and round 11 (green), with yellow indicating areas of overlap. (b) Quantification of displacement. A region with very low density was selected in a sample probe for Wfs1 mRNA, allowing each focus in round 1 (blue cross) to be registered with its corresponding focus in round 11 (yellow cross). Center of each focus was determined using find maxima function in Fiji with a prominence of 180. The plot on the right displays the distances (in nm) between corresponding foci, with the X and Y axis indicates the position in µm of each foci and the values on the plot represent the displacement in nm at a resolution of 300nm. The average displacement was calculated by fitting the binned values of displacement (bins = 0-300, 300-400,400-500) to a continuous normal distribution. (c) Number of foci across all rounds of hybridization/stripping of fluorescent probes from the area selected in (a). Note: All images were initially aligned on the dapi signal.

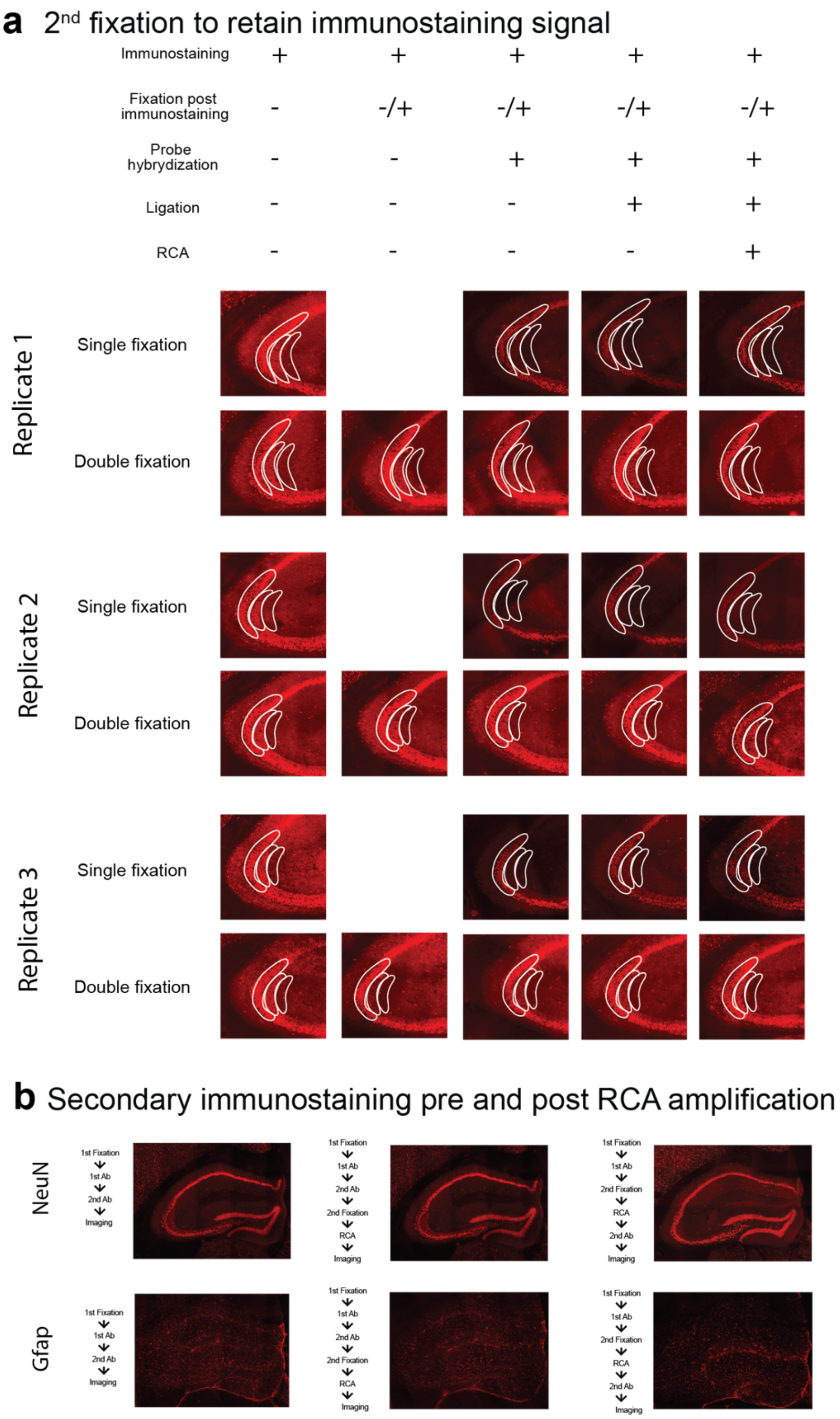


**Supplementary Fig. 2. Effect of post-fixation on conventional immunostaining.** (a) Microscope image of conventional NeuN immunostaining under different conditions, with corresponding conditions listed above. The white lines outline the areas from all replicates used for calculating the average signal presented in the bar chart shown in Fig. 1b. (b) Microscope image of NeuN (top) and GFAP (bottom) immunostainings comparing conditions without Cassini (left), with secondary immunostaining before 2nd fixation and RCA (center) and after 2nd fixation and RCA (right).

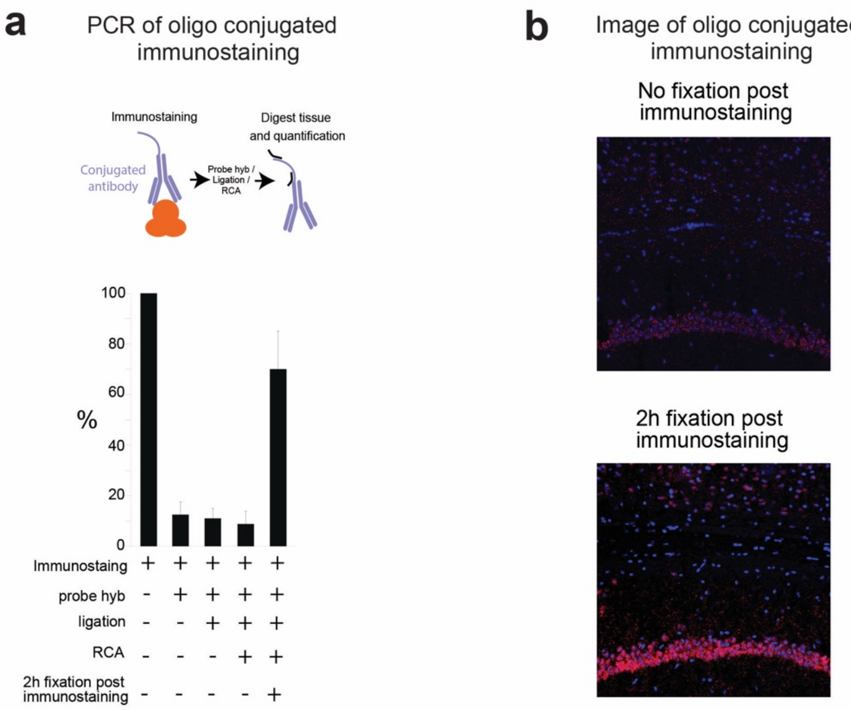


**Supplementary Fig. 3. Effect of Cassini and post-fixation on oligo-conjugated immunostaining.** (a) The total amount of conjugated anti-NeuN antibody was evaluated by quantifying the PCR amplification of the conjugated oligo after digestion of the brain hemisphere. (b) Microscopy image with DAPI signal in blue and immunostaining signal in red (bottom).

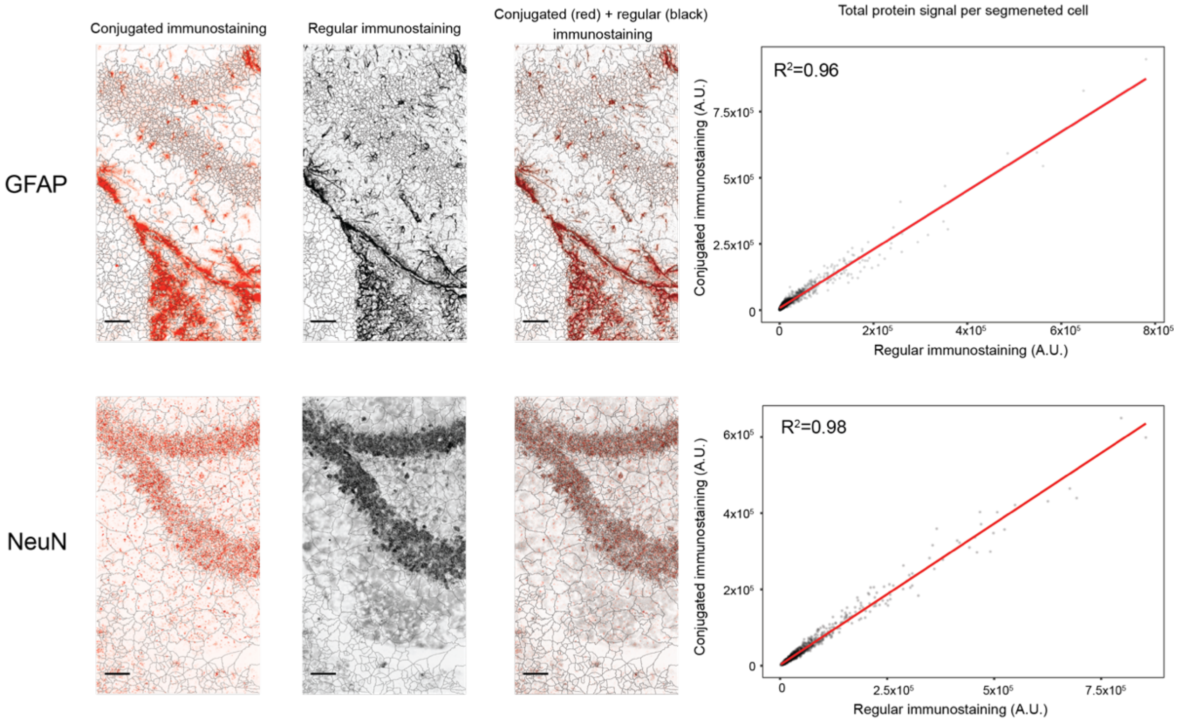


**Supplementary Fig. 4. Correlation of conventional and oligo-conjugated immunostainings.** The microscope images were segmented based on the DAPI signal. The total protein signal within each cell was calculated for both immunostainings. Scale bar at the bottom left of the microscope images = 100 μm.

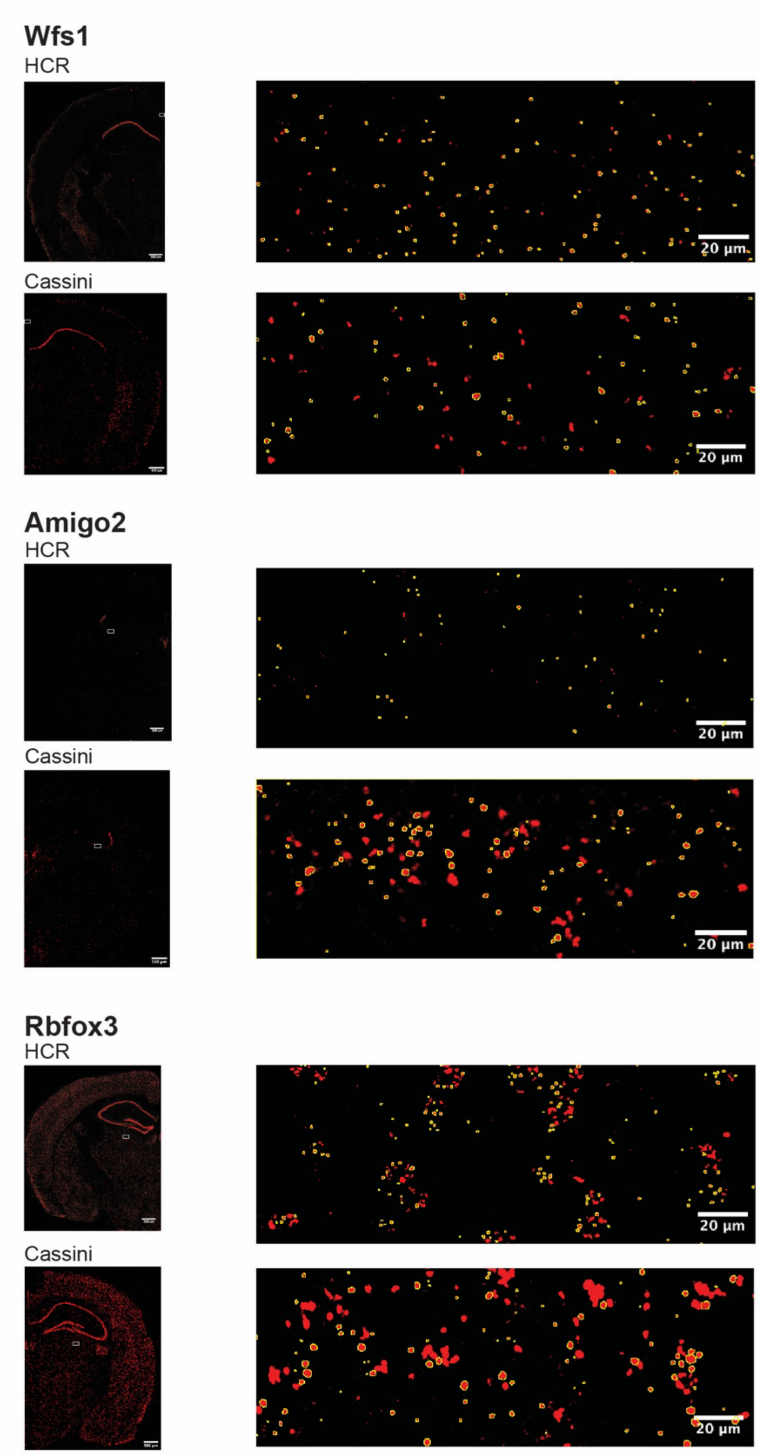


**Supplementary Fig. 5. Foci size comparison between Cassini and HCR**. On the left, the half hemisphere with a white rectangle indicating the zoomed area shown on the right. To facilitate selection of single dots, area with sparse foci were selected and particles with circularity >0.85 and area >0.3 $\mu m^2$ were outlined.

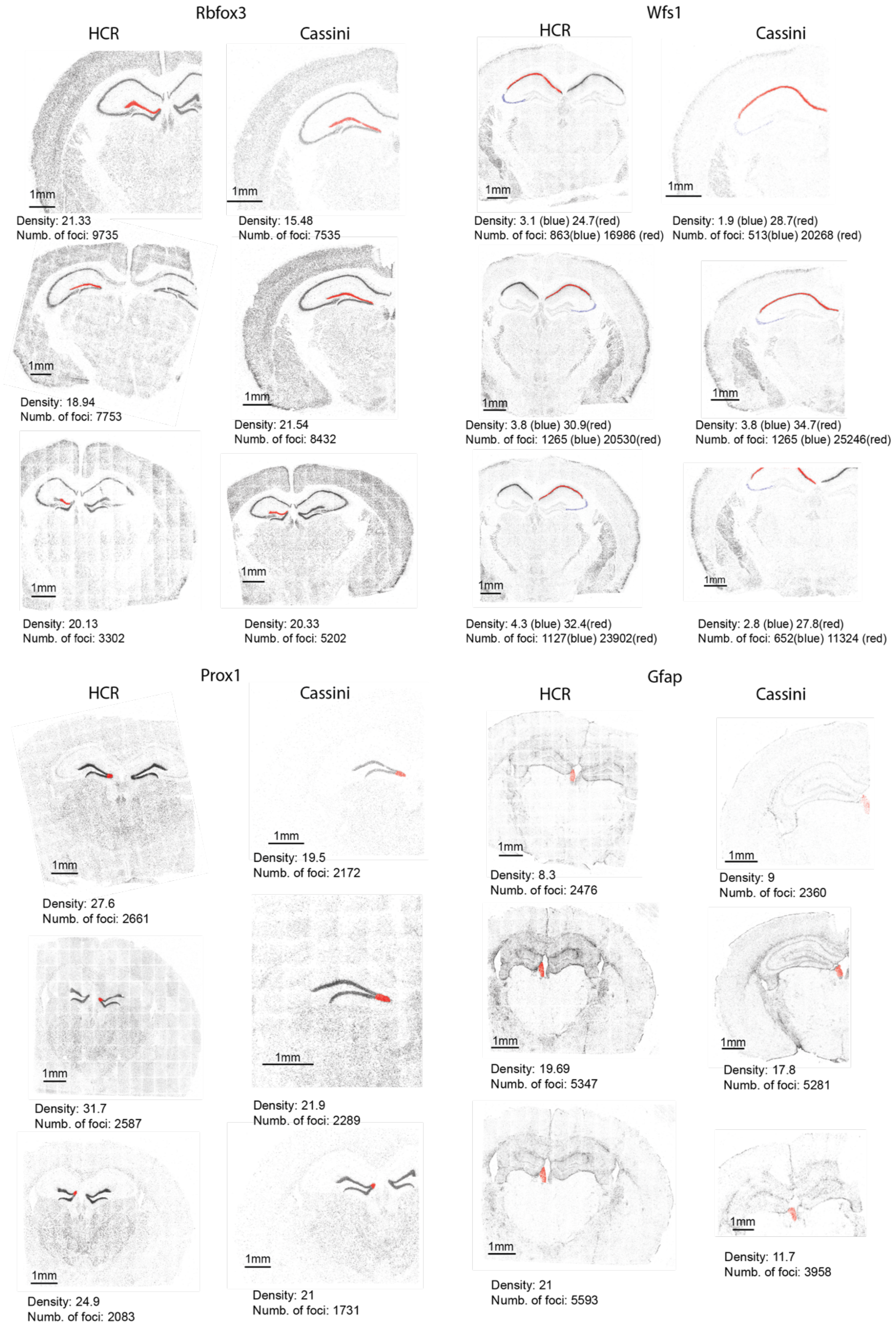
Rbfox3
HCR
Cassini
1mm
Density: 21.33
Numb. of foci: 9735
Density: 15.48
Numb. of foci: 7535
Density: 18.94
Numb. of foci: 7753
Density: 21.54
Numb. of foci: 8432
Density: 20.13
Numb. of foci: 3302
Density: 20.33
Numb. of foci: 5202
Wfs1
HCR
Cassini
Density: 3.1 (blue) 24.7(red)
Numb. of foci: 863(blue) 16986 (red)
Density: 1.9 (blue) 28.7(red)
Numb. of foci: 513(blue) 20268 (red)
Density: 3.8 (blue) 30.9(red)
Numb. of foci: 1265 (blue) 20530(red)
Density: 3.8 (blue) 34.7(red)
Numb. of foci: 1265 (blue) 25246(red)
Density: 4.3 (blue) 32.4(red)
Numb. of foci: 1127(blue) 23902(red)
Density: 2.8 (blue) 27.8(red)
Numb. of foci: 652(blue) 11324 (red)
Prox1
HCR
Cassini
Density: 19.5
Numb. of foci: 2172
Density: 27.6
Numb. of foci: 2661
Density: 31.7
Numb. of foci: 2587
Density: 21.9
Numb. of foci: 2289
Density: 24.9
Numb. of foci: 2083
Density: 21
Numb. of foci: 1731
Gfap
HCR
Cassini
Density: 8.3
Numb. of foci: 2476
Density: 9
Numb. of foci: 2360
Density: 19.69
Numb. of foci: 5347
Density: 17.8
Numb. of foci: 5281
Density: 21
Numb. of foci: 5593
Density: 11.7
Numb. of foci: 3958

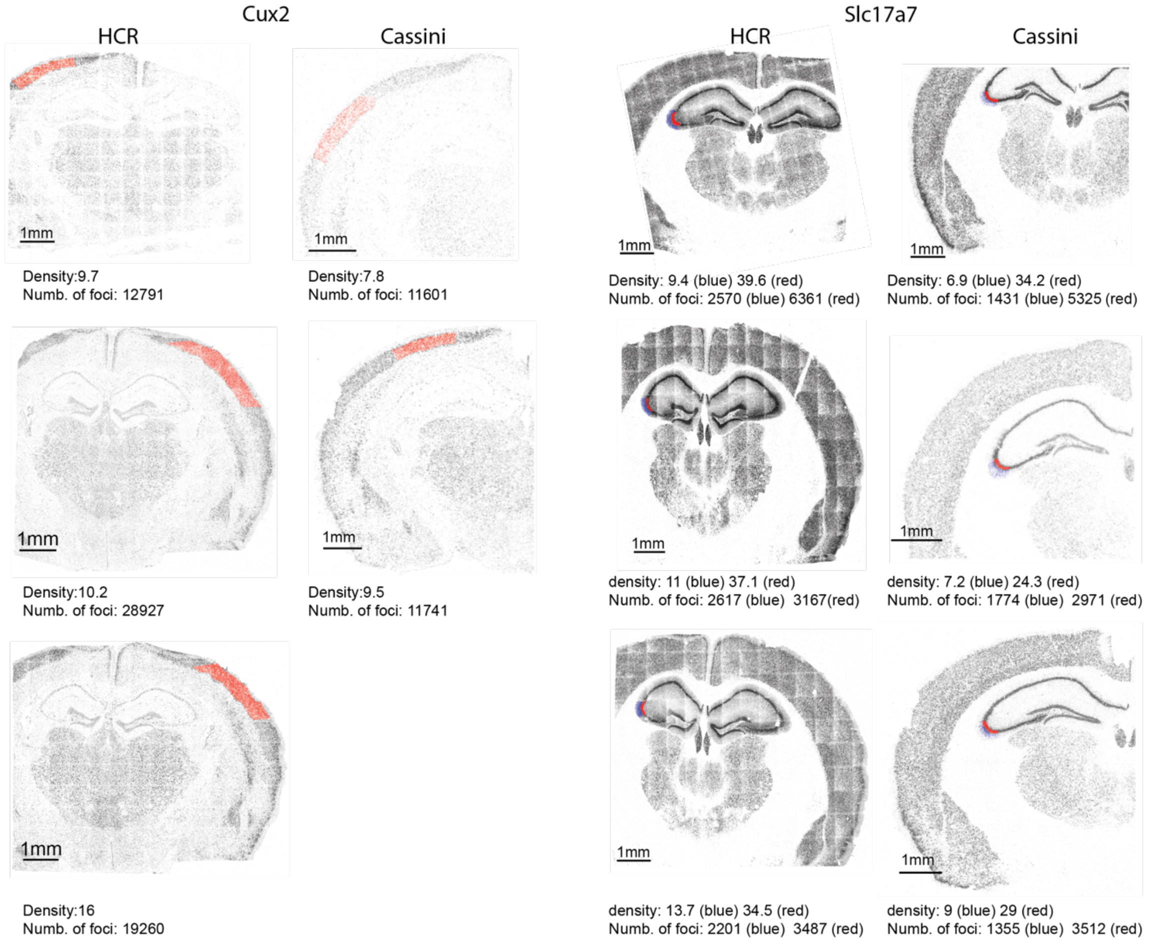


**Supplementary Fig. 6. Comparison of HCR and Cassini mRNAs detection.** Each image shows all foci detected in a brain slice for a specific gene. Density is calculated as follows: An area is selected (red dots), and for each dot in the area, the number of neighboring foci within a radius of 10 µm is determined.

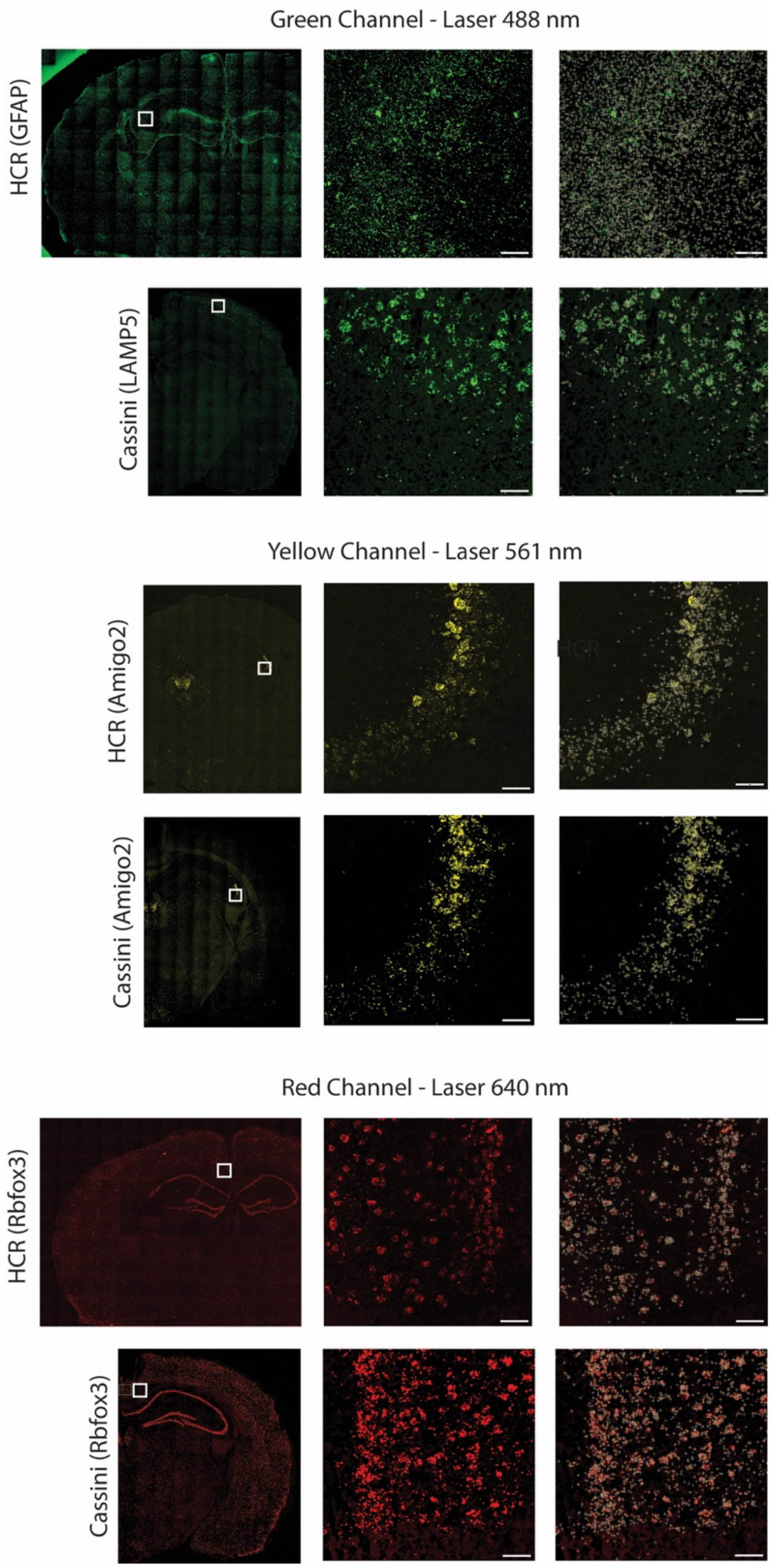


**Supplementary Fig. 7. Foci detection using the *Find Maxima* function in FIJI**. On the left are the full tissue sections stained for the gene indicated on the side, with a white square marking the zoomed area. The center and the right images are the zoom area with and without foci detection marks (white crosses with yellow central dots). The scale bar at the bottom right of the zoomed images represents 40 µm. The prominence parameters are set to 200 for the Cassini images and 40 for the HCR images

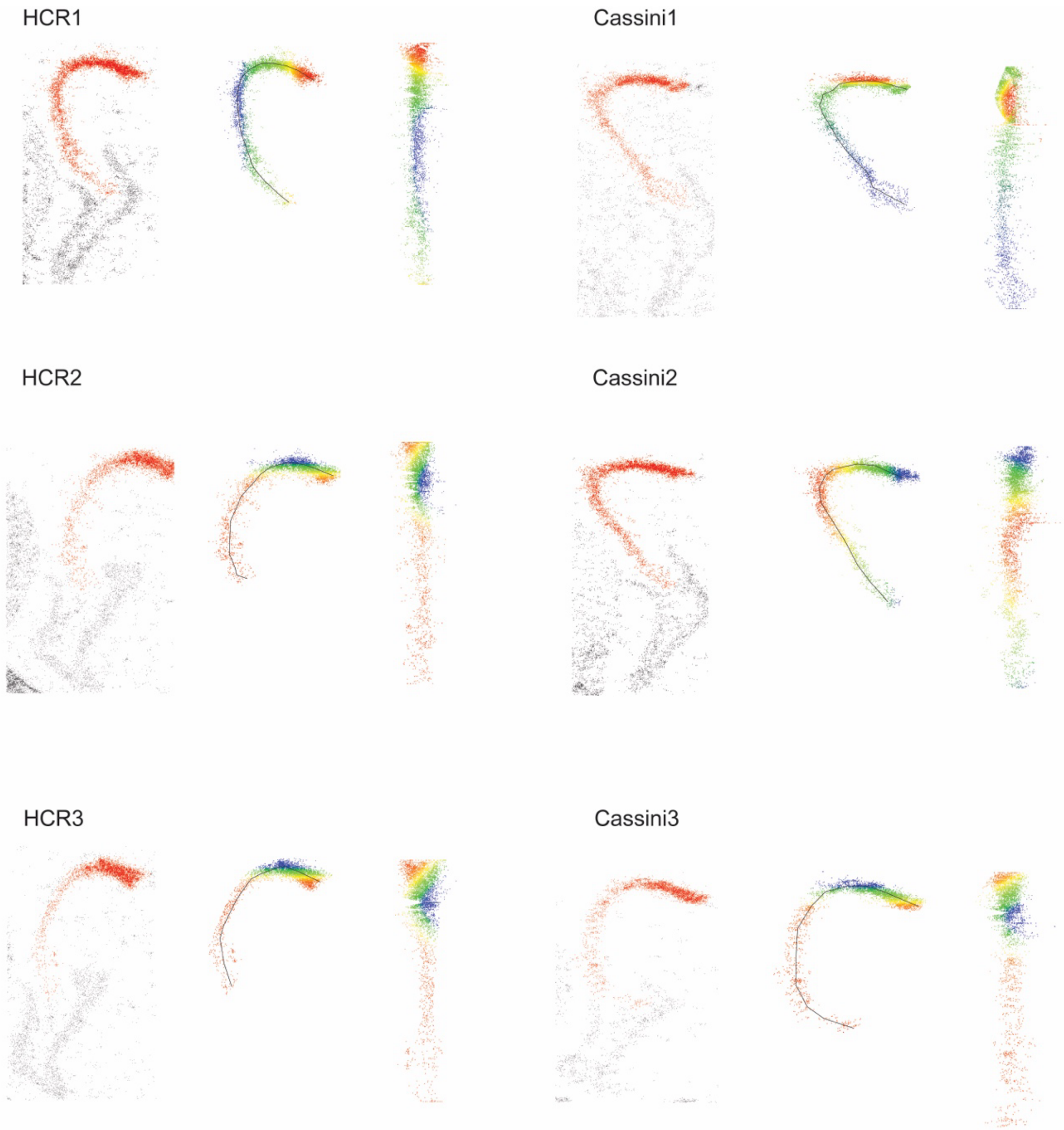


**Supplementary Fig. 8**. **Gradient of Amigo2 in CA2/CA3 area**. The CA2/CA3 area and the line of curvature were both outlined manually, and the data were straightened along the line of curvature. Rainbow colors are used to help visualize the straightening process.

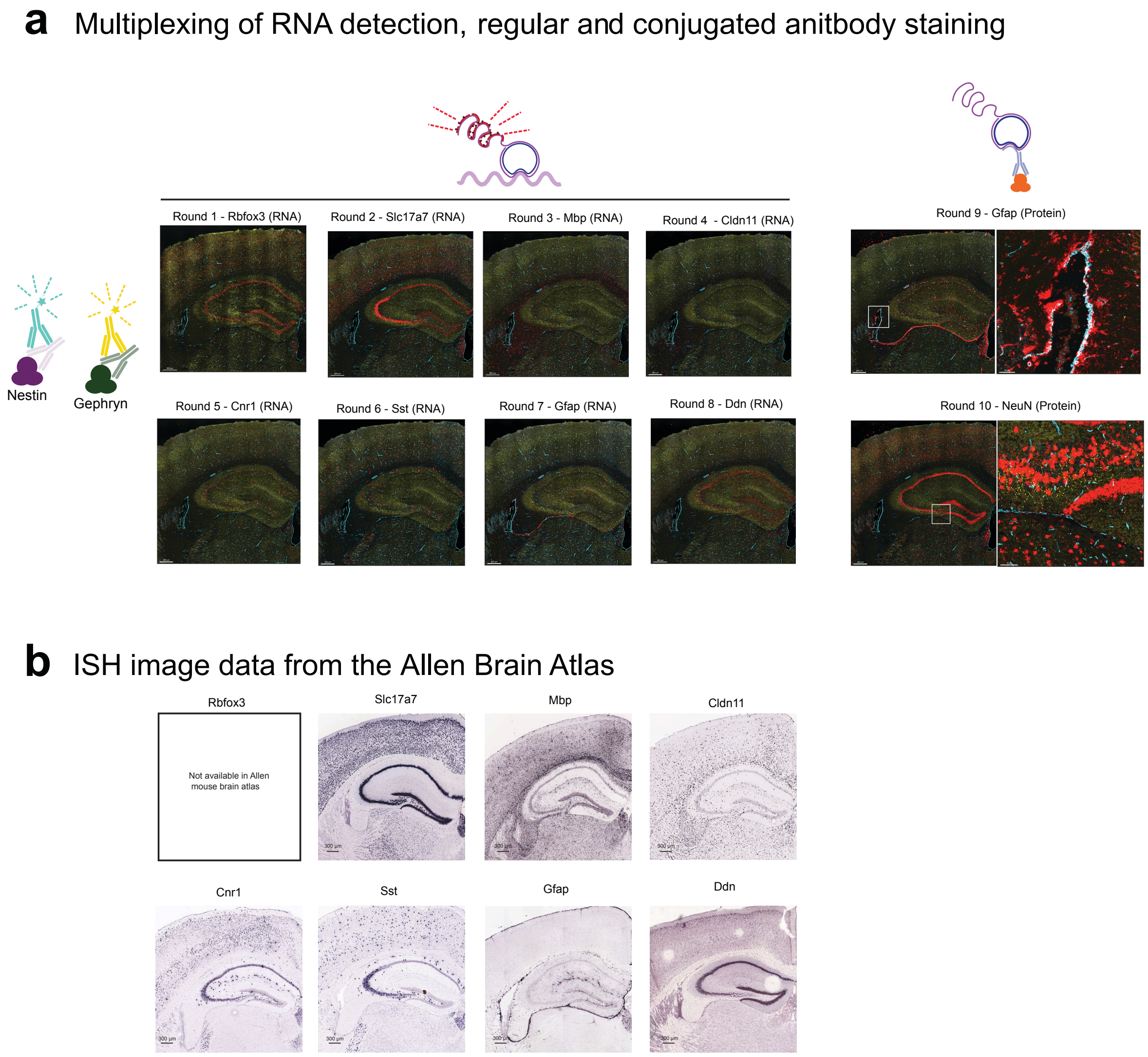


**Supplementary Fig. 9** (a) Microscope images of 10-round Cassini to survey 8 RNA types, 2 conventional and 2 oligo conjugated immunostainings. In each round two fluorescent channels are used to show immunostaining of Nestin (Yellow) and Gephryn (turquoise) using conventional antibodies. The third fluorescent channel (red) cycles through the detection of 8 RNA types and 2 oligo-conjugated antibodies (NeuN and GFAP). (b) In situ hybridization (ISH) image from the Allen Brain Atlas (https://mouse.brain-map.org/) showing the genes displayed in (a).

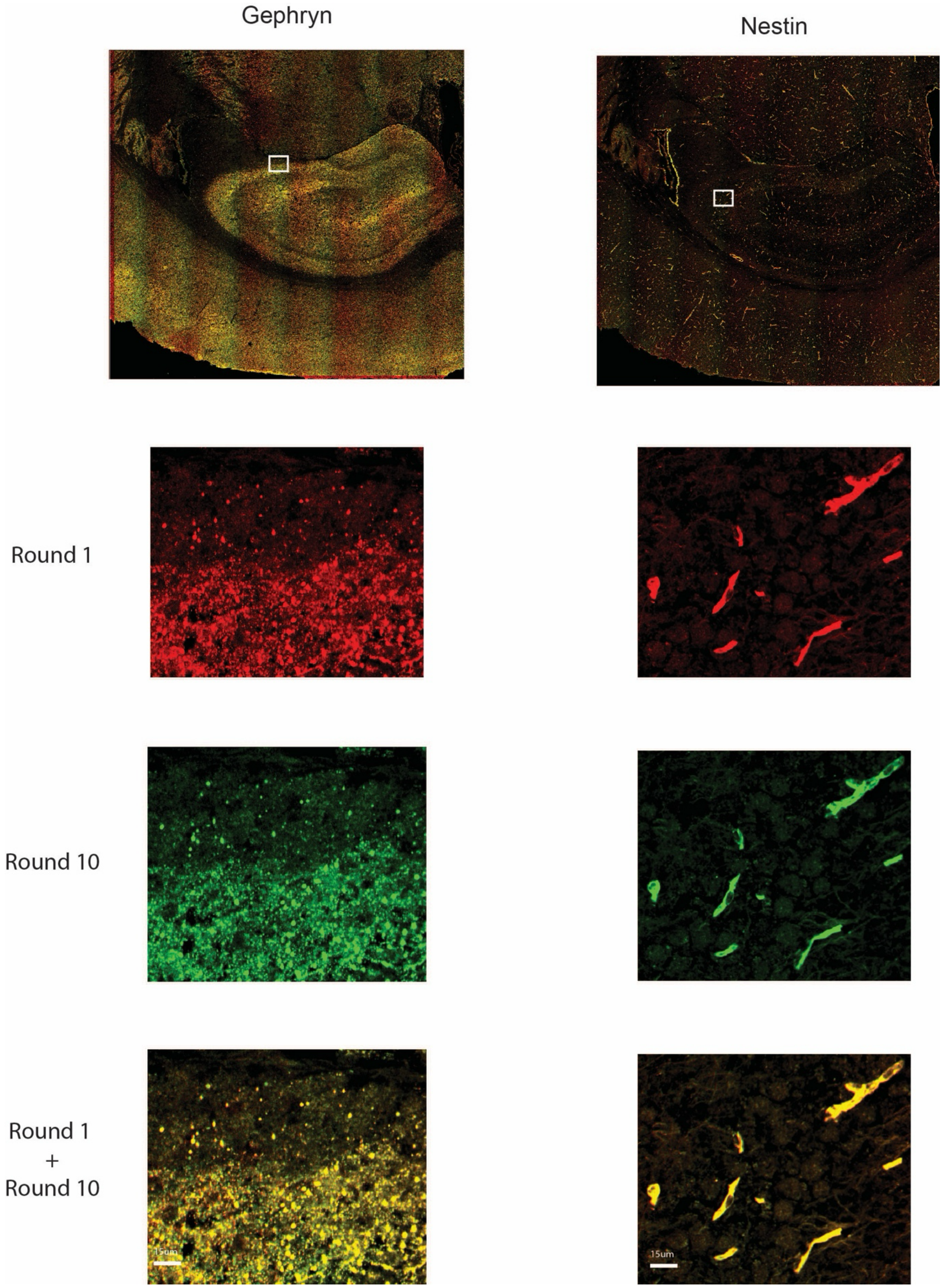


**Supplementary Fig. 10**. Evaluation of the displacement of post-fixed conventional immunostaining after multiple rounds of stripping and hybridization.

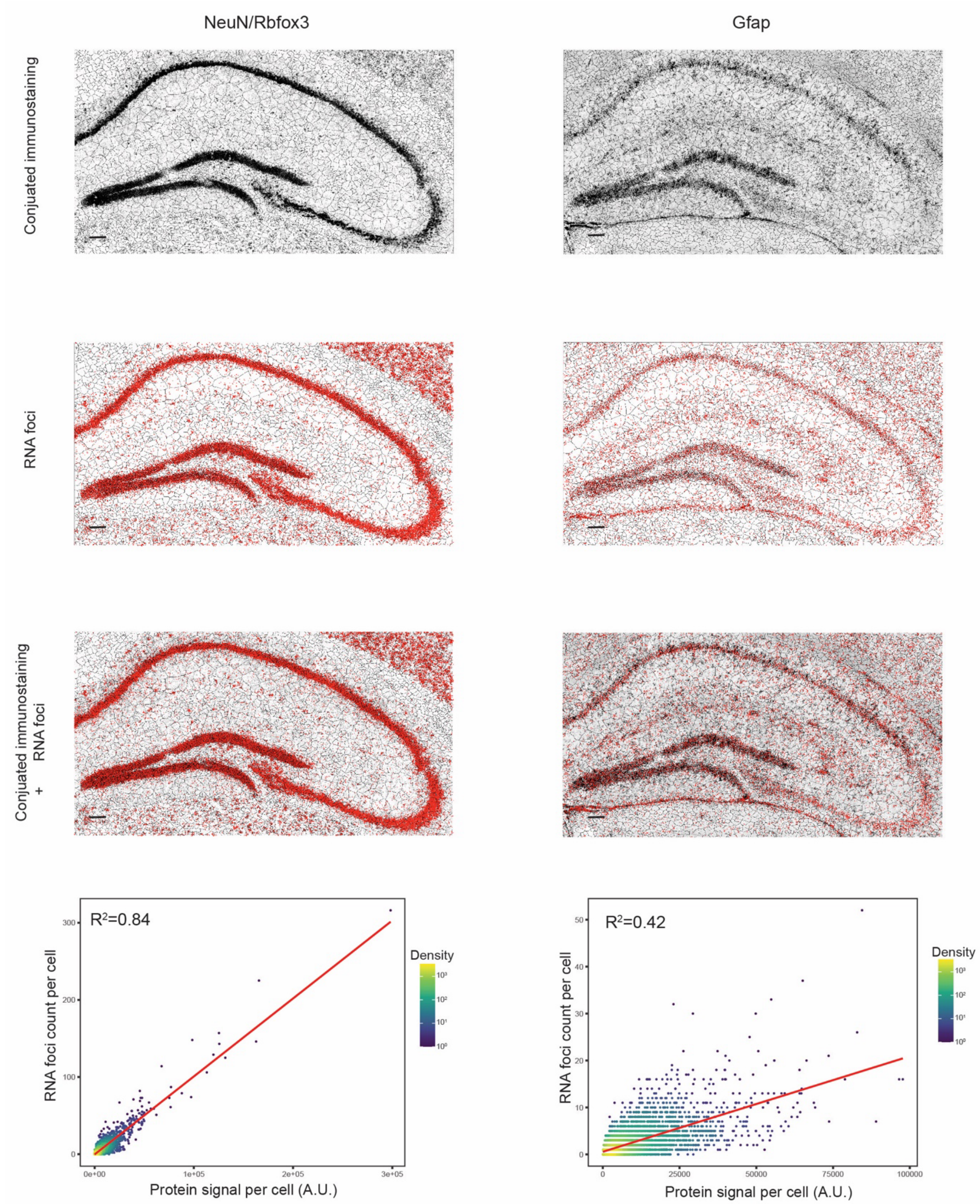


**Supplementary Fig. 11**. **Correlation between RNA count and protein signal.** The microscope images were segmented based on the DAPI signal. Within each segmented cell, the total intensity of immunostaining signals was measured, and the total count of mRNA foci was quantified.

The scale bar at the bottom left of the microscope images = 100 μm

### Detailed workflow and protocol

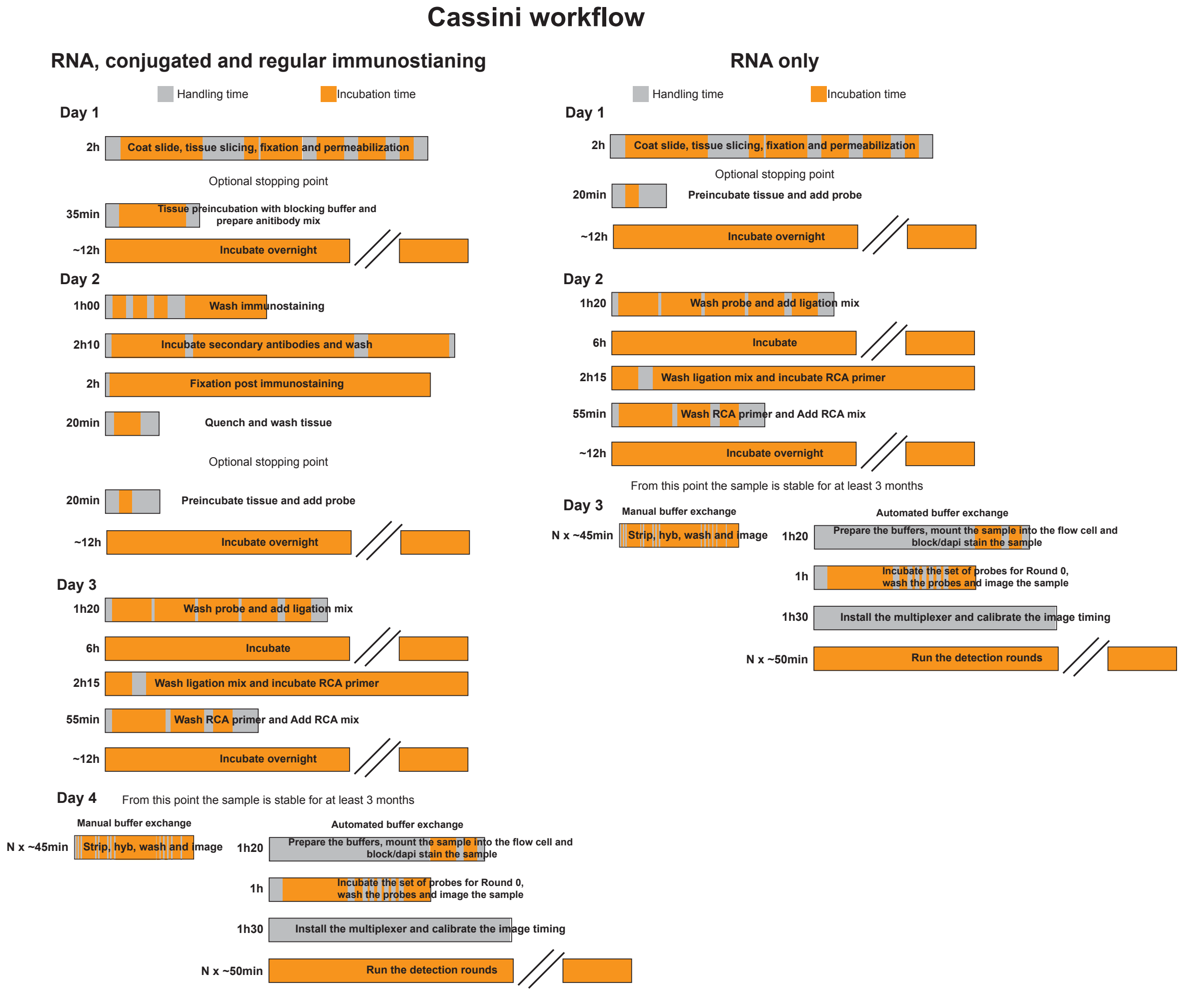


**Note: The time estimate is calculated considering that the mixes are prepared during the incubation period, making some incubation time bar length shorter than the full incubation time**

**Common reagents for RNA and immunostaining**

- Padlock probes : **Padlock probes need to be phosphorylated on the 5'**
- Fluorescent probes
- RCA primer- **RCA primer needs 2 phosphorothioate bonds on the 3' to prevent degradation – e.g. CCTGTGTGAGTCTCC*T*G**
- PBS
- Ultra pure Water
- Tris-HCL pH7.5 1M
- HCl 1M
- Poly-L-Lysine Solution (0.01%) (Sigma A-005-C)
- 40mm glass coverslips (Bioptechs Inc)
- 9x9mm hybridization chamber (Biorad SLF0201)
- 4% PFA (Thermo Fisher Scientific J61899.AK)
- Triton X (Sigma X100-5ML)
- RNAse inhibitor (Y9240L Qiagen)
- Formamide (Sigma Aldrich F7503-4L)
- 20XSSC (Thomas Scientific C000A15)
- Dapi (Thermo Fisher 62248)
- High molecular weight dextran sulfate (Sigma Aldrich S4031)
- FcR Blocking Reagent mouse (Miltenyi biotec 130-092-575)
- Ultra pure BSA 5% (Invitrogen 56773)
- SplintR Ligase (NEB M0375L)
- Phi29 (Thermo Fisher EP0094)
- dNTPs (NEB N0447L)

**Reagents for immunostaining (regular + conjugated)**

- FcR Blocking Reagent mouse (Miltenyi biotec #130-092-575)
- Ultra pure BSA 5% (Invitrogen #56773)

**Reagents for conjugated immunostaining**

- Dextran sulfate-4K (Sigma #75027)
- NaCl 1M
- oYo-Link® Oligo Custom (AlphaThera)

**Reagents for regular immunostaining**

- Normal serum specific to the species of the secondary

**Hardware Material**

Sample preparation (Optional)

- HybEZ™ II Oven (ACDBio)

Automated buffer exchange (Optional)

- FCS2 flow cell (Bioptechs)
- 2 Rotary valves (AMF)
- Raspberry pi
- Peristaltic pump (Kamoer KCM-ODM-B253)

#### Tissue preparation

##### Material
- Ultra pure Water
- Poly-L-Lysine Solution (0.01%) (Sigma A-005-C)
- 40mm glass coverslips (Bioptechs Inc)
- 9x9mm hybridization chamber (Biorad SLF0201)

##### Protocol
1. Coat the round glass with a solution of Poly-L-Lysine Solution (0.01%) for 30 minutes. Wash them by immersion in Ultra pure H2O and let them dry until all moisture has evaporated. If you want to speed up the drying process, you may use compressed air. During the incubation time, place the fresh frozen brain in the cryostat at -18˚C to equilibrate (at least 20 minutes).

2. Add a hybridization chamber on each slides (Biorad) and press firmly with a plastic tool
   Critical step: The glass is very thin and can break easily if press on an uneven surface

3. Place the glass slides in the cryostat at -18˚C and wait for 5 minutes before slicing. Slice a 10um thick tissue section and melt it on the glass slide with the help of a finger. Let the slide equilibrate at RT from 2 minutes in a humidified chamber before processing to the next step.

#### Tissue fixation

##### Material
- PBS
- Ultra pure Water
- 4% PFA (Thermo Fisher Scientific J61899.AK)
- Tris-HCL pH7.5 1M
- Triton X (Sigma X100-5ML)
- HCl 1M

##### Mixes
Tris-HCL pH7.5 100mM (Solution is stable at room temp for long term storage)
- 900 µl of ultra pure water
- 100 µl Tris-HCL pH 7.5 1M

Triton 0.25% (Solution is stable at room temp for long term storage)
- 10 ml of Ultra pure water
- 25 µl of Triton X → To facilitate easier pipetting of the Triton solution, we recommend trimming the ends of the pipette tips.

HCl 100mM (Solution is stable at room temp for long term storage)
- 900 µl of Ultra pure Water
- 100 µl of HCl 1M

**Protocol**

4. Add 50 µl of PFA 4% and incubate for 12-14 minutes at room temperature and wash 3 times with 80 µl of PBS

5. Add 50 µl of 20mM Tris-HCL to quench the PFA and incubate 10min at room temperature and wash 3 times with 80 µl of PBS

6. Add 30 µl of 0.25% Triton and incubate 10min at room temperature and wash 3 times with 80 µl of PBS

7. Add 50 µl of 0.1M HCl and incubate 5min at room temperature and wash 3 times with 80 µl of PBS

#### <u>Immunostaining + 2nd fixation (Skip this step if RNA only)</u>

**Material**

- PBS
- Ultra pure Water
- NaCl 1M
- Low molecular weight Dextran sulfate (Sigma Aldrich 75027)
- FcR Blocking Reagent mouse (Miltenyi biotec 130-092-575)
- Ultra pure BSA 5% (Invitrogen 56773)
- RNAse inhibitor (Y9240L Qiagen)
- Tris-HCl pH 7.5 100mM
- Antibodies (conjugated and/or conventional)

**Mixes**

20% Dextran sulfate-4K (Solution is stable at 4˚C for long term storage)
- 1g of Dextran sulfate-4K
- 5ml of ultra pure water

Blocking buffer 1 – Make it fresh
- 1 µl of FcR Blocking Reagent
- 5 µl of NaCl 1M
- 1 µl of RNAse inhibitor
- 10 µl of Ultra pure BSA 5%
- 5 µl of 20% Dextran sulfate-4K
- 78 µl of PBS

Blocking buffer 2 – Make it fresh
- 2.5 µl of FcR Blocking Reagent
- 12.5 µl of NaCl 1M
- 25 µl of Ultra pure BSA 5%
- 12.5 µl of 20% Dextran sulfate-4K
- 197.5 µl of PBS

PBS + RNAse inhibitor – Make it fresh
- 160 µl of PBS
- 1.6 µl of RNAse inhibitor

Secondary blocking buffer – Make it fresh (Only for conventional immunostaining)
  - 5 µl of Normal serum specific to the species of the secondary
  - 94 µl of PBS
  - 1 µl of RNAse inhibitor

**Protocol**
**All steps must be done in a humidified chamber !**

8. Add 50 µl of blocking buffer 1 and incubate 30min at 4c.

9. Prepare a diluted solution of antibodies by incorporating them into the blocking buffer. To achieve optimal specificity, it is generally necessary to dilute the conjugated antibodies 100 to 200 more than standard antibodies. We typically dilute our conjugated antibodies to 100’000x. Remove previous solution and add 50 µl of the diluted antibodies to each sample and incubate overnight at 4C.

10. Wash 3 times with 80 µl of blocking buffer 2, with 5 min incubation at room temp for each wash.

11. Wash 3 times with 80 µl PBS with ~15 seconds of incubation for each wash and add 80 µl of PBS + RNAse inhibitor (0.4U/ µl) and incubate 30 min at room temp.

12. *Only for conventional immunostaining*: Incubate the sample with 50µl Secondary blocking buffer for 30min at room temperature. Dilute secondary antibodies in 50µl of secondary blocking buffer, remove the blocking solution, add the secondary mix to the sample and incubate 1h at room temp.

13. *Only for conventional immunostaining*: Wash the secondary antibodies 3 times with 80µl of PBS with ~15 seconds of incubation for each wash and incubate with 80µl PBS + RNAs inhibitor for 30 minutes at room temperature. Wash with 80ul of PBS.

14. Remove PBS and add 50 µl of PFA 4% and incubate for 2h at room temperature.

15. Remove the PFA and add 50 µl of 100mM Tris-HCL and incubate 10min at room temperature.

16. Wash 3 times with 80 µl of PBS, with 1 min incubation at room temp for each wash.

#### Probe hybridization

**Material**
- Ultra pure Water
- Formamide (Sigma Aldrich F7503-4L)
- 20X SSC (Thomas Scientific C000A15)

- RNAse inhibitor (Y9240L Qiagen)
- Padlock probes

**Mixes**

Wash-20 (Solution is stable for a week)

- 2ml of formamide
- 1ml of SSC 20x
- 7ml of ultra pure water

Wash-20 + Rnase inhibitor – Make it fresh

- 50 µl of Wash-20
- 0.5 µl of RNAse inhibitor

Padlock probe solution – Make it fresh

- 3 µl of 20X SSC
- 6 µl of formamide
- 0.6 µl of RNAse inhibitor
- 0.6 µl of Padlock probe 500nM
- 19.8 µl of $H_2O$

**Protocol**

**All steps must be done in a humidified chamber!**

17. Add 50 µl of Wash-20 + RNAse inhibitor and incubate for 15min at room temperature.

18. Prepare the 30 µl Padlock probe solution to a final concentration of 10nM. Incubate the solution overnight at 37˚C.

#### Probe wash and ligation

**Material**

- PBS
- Wash-20
- SplintR ligase kit (NEB M0375L)
- RNAse inhibitor (Y9240L Qiagen)

**Mixes**

Wash-20 + Rnase inhibitor – Make it fresh

- 150 µl of Wash-20
- 1.5 µl of RNAse inhibitor

Ligase mix – Make it fresh and prepare on ice

- 2.5 µl of splintR ligase
- 5 µl of SplintR ligase buffer 10x
- 1 µl of RNAse inhibitor
- 41.5 µl Ultra pure water

**Protocol**

**All steps must be done in a humidified chamber!**

19. Wash 3 times with 50 µl of Wash-20 supplemented + RNAse. Incubate each wash 15min at 37˚C.

20. Wash with 80 µl of PBS and incubate 15min at 37˚C.

21. Wash with 80 µl of splintR ligase buffer 1X and incubate 15min at room temperature.

22. Prepare the ligase mix on ice, remove preincubation buffer, add 50 µl to the sample and incubate for 6h to overnight at 37˚C.

#### RCA

##### Material

- RCA primer
- Phi29 kit (Thermo Fisher Scientific EP0094)
- RNAse inhibitor (Y9240L Qiagen)
- Ultra pure Water
- PBS
- dNTPs 10mM (NEB N0447L)

##### Mixes

RCA primer hybridization mix – Make it fresh

- 200ul of Wash-20
- 1ul of RCA primer (100µM)

RCA mix -Make it fresh and prepare on ice

- 5 µl of Phi29 buffer 10x
- 1.25 µl of 10mM dNTPs
- 4 µl of Phi29 (10U/µl)
- 0.5 µl of RNAse inhibitor
- 39.25 µl of ultra pure Water

##### Protocol

**All steps must be done in a humidified chamber!**

23. Remove the ligation mix and add 50ul Wash-20 buffer for 10 min at RT.

24. Remove Wash-20 preincubation and add of 50 µl of RCA primer hybridization mix and incubate for 2hr at 37˚C.

25. Remove the primer mix and add 80 µl of Wash-20 incubated for 20min at 37˚C.

26. Remove the wash and add 80 µl of PBS incubated for 15min at 37˚C.

27. Remove the PBS and add 80 µl of 1X Phi-29 buffer incubated for 15min at RT. Switch the incubator to 30˚C.

28. Prepare the RCA mix on ice, remove preincubation buffer, add 50 µl to the sample and incubate overnight at 30˚C.

29. Remove RCA mix and add 80 µl of PBS to terminate the reaction. From this point the samples is stable for at least 3 months at 4 ˚C (probably more but we did not test it).

#### Probe detection through manual fluidic exchange

##### Material

- Ultra pure Water
- Formamide (Sigma Aldrich F7503-4L)
- 20X SSC (Thomas Scientific C000A15)
- Dapi Sigma Aldrich S40
- HW dextran sulfate 50% (Sigma Aldrich S4031)
- Wash-20
- Fluorescent probes

##### Mixes per N round

Wash-20-HWDS-Dapi

- N x 300 µl of Wash-20
- N x 0.15 µl of Dapi
- N x 3 µl of HW dextran sulfate 50%

Stripping solution

- N x 400 µl of formamide
- N x 12.5 µl of 20X SSC
- N x 87.5 µl of Ultra pure $H_2O$

N x Probe solutions

- 30 µl of Wash-20
- 0.15 µl of fluorescent probe A 100 µM
- 0.15 µl of fluorescent probe B 100 µM
- 0.15 µl of fluorescent probe C 100 µM

##### Protocol

30. Prepare Wash-20-HWDS-Dapi, stripping solution and all probe solutions. Protect the Wash-20-HWDS-Dapi and fluorescent probe solutions from light.

31. Add 30 µl of first probe solution on the sample and wait 15min at room temp (longer incubation does not affect the staining neither the probe stripping).

    Remove solution and add 100 µl of Wash-20
    Wait for 1 minutes
    Remove solution and add 100 µl of Wash-20
    Wait for 1 minute
    Remove solution and add 100 µl of Wash-20
    Wait for 1 minute
    Remove solution and add 100 µl of Wash-20
    Wait for 1 minute
    Remove solution and add 150 µl of Wash-20
    Wait for 3 minutes
    Image the sample

32. Follow these steps for each round of detection
    a. Probe stripping
       Remove solution and add 150 μl of stripping buffer
       Wait for 30 seconds
       Remove solution and add 150 μl of stripping buffer
       Wait for 30 seconds
       Remove solution and add 150 μl of stripping buffer
       Wait for 5 minutes
    b. Tissue blocking and dapi
       Remove solution and add 150 μl of Wash-20-HWDS-Dapi
       Wait for 30 seconds
       Remove solution and add 150 μl of Wash-20-HWDS-Dapi
       Wait for 3 minutes
    c. Pre incubation
       Remove solution and add 150 μl of Wash-20
       wait for 30 seconds
       Remove solution and add 150 μl of Wash-20
       Wait for 1 minute
    d. Probe incubation
       Remove solution and add 30 μl of probe solution
       Wait for 15 minutes
    e. Probe wash
       Remove solution and add add 150 μl of Wash-20
       Wait for 1 minutes
       Remove solution and add 150 μl of Wash-20
       Wait for 1 minute
       Remove solution and add 150 μl of Wash-20
       Wait for 1 minute
       Remove solution and add 150 μl of Wash-20
       Wait for 1 minute
       Remove solution and add 150 μl of Wash-20
       Wait for 3 minutes
    f. Imaging
       Variable time

#### Probe detection through automated fluidic exchange

**Material**
- Ultra pure Water
- Formamide (Sigma Aldrich F7503-4L)
- 20X SSC (Thomas Scientific C000A15)
- Dapi Sigma Aldrich S40
- HW dextran sulfate 50% (Sigma Aldrich S4031)
- Fluorescent probes

**Mixes**

Wash-20
- 50 ml of formamide
- 25 ml of 20X SSC

- 175 ml of ultra pure water

Wash-20-HWDS-Dapi
- 65 ml of Wash-20
- 16.25 µl of Dapi
- 650 µl of HW dextran sulfate 50%

Stripping solution
- 56 ml of formamide
- 1.75 ml of 20X SSC
- 12.25 ml of Ultra pure H2O

N x Probe solutions
- 1 ml of Wash-20
- 3 µl of fluorescent probe A 100 µM
- 3 µl of fluorescent probe B 100 µM
- 3 µl of fluorescent probe C 100 µM

**Protocol**

30 Prepare Wash-20, Wash-20-HWDS-Dapi, stripping solution and all probe solutions. Protect the Wash-20-HWDS-Dapi and fluorescent probe solutions from light.

31 ▲CAUTION Remove the hydrophobic barrier using a sharp blade (the glass slide is very fragile and can break if not handle correctly). Place carefully the sample in the FCS2 flow cell and close the flow cell according to the manufacturer instructions. Connect separate tubing to each outlet of the flow cell and introduce 1ml of Wash20-DSS-Dapi with a syringe. Immediately close the tubing circuit, by connecting the same tubing on both side of the flow cell and incubate the solution for 10 minutes at room temperature.

32 introduce 3ml of Wash-20 and incubate for 5minutes at room temperature. After incubation, flow 3ml of Wash-20 and close the tubing of the flow cell, to maintain Wash-20 in the flow cell.

33 Optional: You can manually add the first probe set to the sample

Add 500 µl of probe solution into the flow cell and incubate for at least 25min at room temp (longer incubation does not affect the staining neither the probe stripping).
Add 2ml of Wash-20
Wait for 3 minutes
Add 1ml of Wash-20
Wait for 1 minute
Add 1ml of Wash-20
Wait for 1 minute
Add 1ml of Wash-20
Wait for 1 minute
Add 1ml of Wash-20
Wait for 1 minute

Add 2ml of Wash-20
Wait for 1 minute

34 Connect all tubing from the valve controller to the probe, blocking, wash and stripping solutions. The first valve controller receives the probe mixes (one mix per cycle), and its outlet is connected to the inlet of the second rotary valve. Additionally, the inlets of the second controller valve are connected to the stripping solution, Wash-20-HWDS-Dapi and wash-20. The outlet of the second rotary valves is connected to the flow cell. The peristaltic pump is placed downstream of the flow cell and controls the flow rate at 0.3 ml/min for the probe mix buffer and 1 ml/min for the other buffers.
To prevent the formation of bubbles, we recommend plugging tubes into all empty ports on the valve controller and connect those tubes to Wash-20.

35 Prime all tubes in this order:
a. 200 µl of fluorescent probe tubes
b. 200 µl of all filler tubes plug on the empty ports of the valve controller.
c. 1ml of stripping buffer
d. 1ml of Wash-20-HWDS-Dapi
e. 2ml of Wash-20

Once all tubes are primed, visually inspect that all tubes are bubble free, and connect the outlet of the valve controller to the flow cell. Flow 1ml of Wash-20 and image the round 0 from the manual probe hybridization.

36 Synchronize the automated fluidic exchange with the image acquisition using a mouse autoclicker or time series from the microscope software control.

37 Run the Sequential hybridization with the following program:
Timing ~47minutes per cycles (without imaging time)

a. <u>Probe stripping</u>
2.5ml of stripping buffer
Wait for 30 seconds
1ml of stripping buffer
Wait for 30 seconds
2.5ml of stripping buffer
Wait for 5 minutes
b. <u>Tissue blocking and dapi</u>
2.5ml of Wash-20-HWDS-Dapi
Wait for 30 seconds
2.5ml of Wash-20-HWDS-Dapi
Wait for 3 minutes
c. <u>Pre incubation</u>
2.5ml of Wash-20
wait for 30 seconds
2.5ml of Wash-20
Wait for 2 minutes
d. <u>Probe incubation</u>
600 µl of probe solution

Wait for 25 minutes

e. Probe wash

Add 2ml of Wash-20
Wait for 1 minutes
Add 1ml of Wash-20
Wait for 1 minute
Add 1ml of Wash-20
Wait for 1 minute
Add 1ml of Wash-20
Wait for 1 minute
Add 1ml of Wash-20
Wait for 1 minute
Add 2ml of Wash-20
Wait for 1 minute

f. Imaging

Variable time
